# Genomic Surveillance of COVID-19 Variants with Language Models and Machine Learning

**DOI:** 10.1101/2021.05.25.445601

**Authors:** Sargun Nagpal, Ridam Pal, Ashima, Ananya Tyagi, Sadhana Tripathi, Aditya Nagori, Saad Ahmad, Hara Prasad Mishra, Rishabh Malhotra, Rintu Kutum, Tavpritesh Sethi

**Author notes:** Contributed equally to this work and share first authorship (Sargun Nagpal, Ridam Pal). Contributed equally and share second authorship (Ashima, Ananya Tyagi, Sadhana Tripathi).

## Abstract

The global efforts to control COVID-19 are threatened by the rapid emergence of novel SARS-CoV-2 variants that may display undesirable characteristics such as immune escape, increased transmissibility or pathogenicity. Early prediction for emergence of new strains with these features is critical for pandemic preparedness. We present *Strainflow*, a supervised and causally predictive model using unsupervised latent space features of SARS-CoV-2 genome sequences. *Strainflow* was trained and validated on 0.9 million sequences for the period December, 2019 to June, 2021 and the frozen model was prospectively validated from July, 2021 to December, 2021. *Strainflow* captured the rise in cases two months ahead of the Delta and Omicron surges in most countries including the prediction of a surge in India as early as beginning of November, 2021. Entropy analysis of *Strainflow* unsupervised embeddings clearly reveals the explore-exploit cycles in genomic feature-space, thus adding interpretability to the deep learning based model. We also conducted codon-level analysis of our model for interpretability and biological validity of our unsupervised features. *Strainflow* application is openly available as an interactive web-application for prospective genomic surveillance of COVID-19 across the globe.

## 1. Introduction

New variants of SARS-CoV-2 continue to rage across the globe causing devastating waves of the pandemic. Such waves may continue to occur and many lives can be saved through early preparedness. COVID-19 is reported to have claimed 5.45 million lives as of Jan 10, 2022 (WHO Coronavirus (COVID-19) Dashboard). A large number of these deaths are attributed to unexpected surges in infections caused by new strains with higher pathogenicity such as the Delta variant of SARS-CoV-2, prompting international health organizations such as the CDC and WHO to declare these as variants of concern (CDC, 2022).The most recent surge of Omicron across the globe with its potential for escaping immunity has seriously undermined the efficacy of global vaccination programs. Most studies around the globe have focussed on forecasting case time series using traditionally reported administrative data. Standard epidemiological approaches such as compartmental and agent-based modeling have been used extensively for forecasting COVID-19 caseloads (Arora et al., 2020). Additionally, numerous studies have used time series analysis, social media mining and multimodal approaches have been utilized for case predictions (Ayan et al., 2021; Kapoor et al., 2020; Melin et al., 2020; Qin et al., 2020; Reiner et al., 2020; Rodríguez et al., 2020; Wu et al., 2020). Earlier, initiatives such as *Nextstrain* (Hadfield et al., 2018) have focused on providing high-quality tracking information for the strains and lineages as these emerge without forecasting or predictions. Hence early prediction of caseloads and emerging variants through genomic signals remains an open challenge for COVID-19.

Unsupervised embeddings have been shown to capture highly nonlinear and contextual relationships (Mikolov et al., 2013). Biological sequences contain a plethora of information that can be exploited for genomic surveillance. However, there is a paucity of studies that explore the use of unsupervised embeddings for machine learning based prediction of surges in infections. In these models, codons (tri-nucleotides, 3-mers) translations represent a natural basis for word representations and have been utilized in the past for learning embedding models for modelling various outcomes such as mutation susceptibility and gene sequence correlations (Yilmaz, 2020) (Wu et al., 2021). Recently, Hie et. al used machine learning along with word embedding techniques to model the semantics and grammar of amino acids corresponding to antigenic change to predict the mutations which might lead to viral escape (Hie et al., 2021). Similarly, Maher et. al predicted emerging mutations of SARS-CoV-2 variants and evaluated biological and neural network based predictors of emerging mutations (Maher et al., 2021). Here, we propose *Strainflow (Figure 1)*, a prospectively validated pipeline with prediction and prospective validation of surges two months ahead of time. Our empirical experiments demonstrate interpretable features based on Entropy of the latent space of SARS-CoV-2 spike region, thus aiding an early warning system for emergence of new variants of concern and case surges.

**Figure 1.**
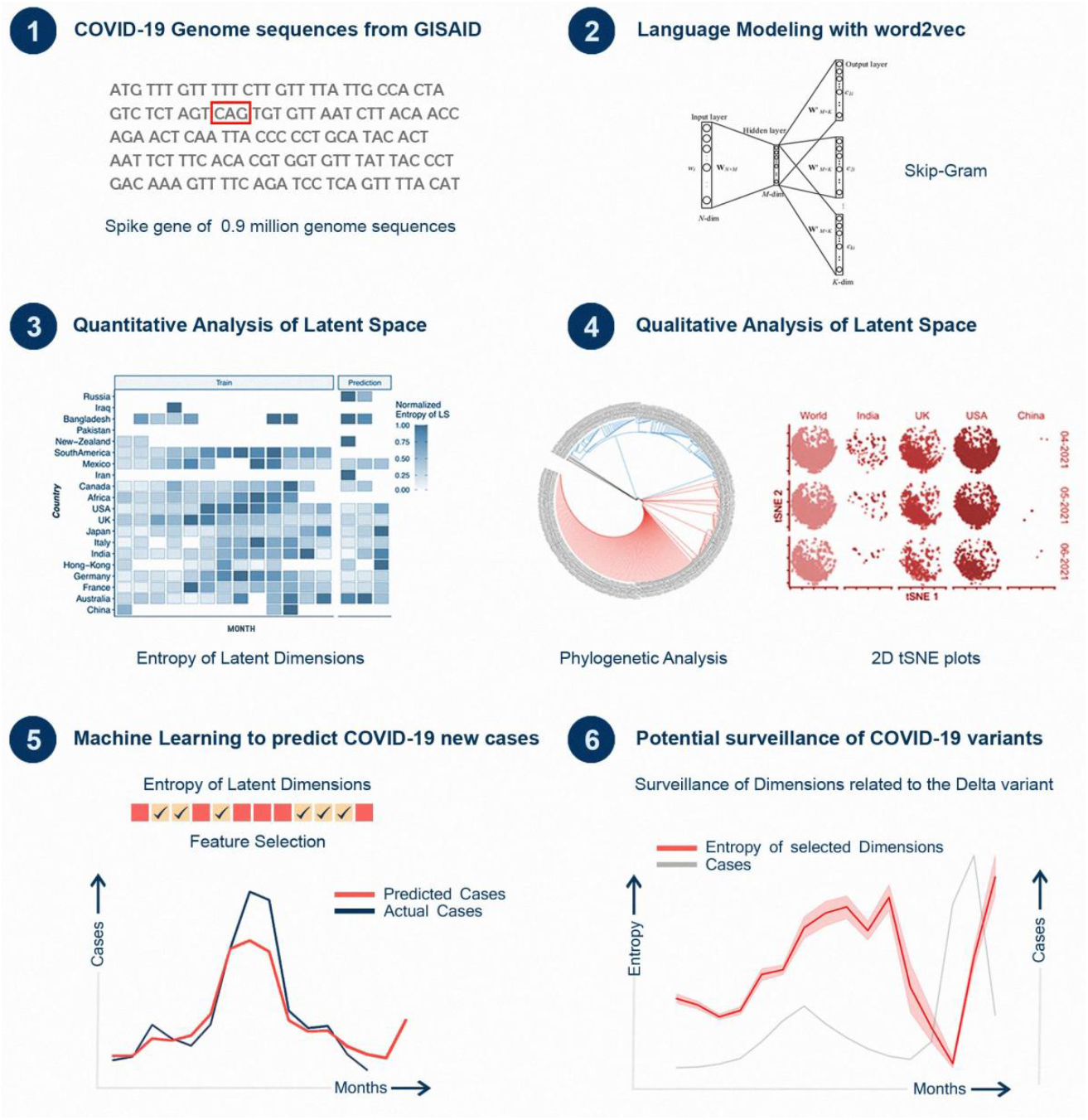
Architecture of the Strainflow pipeline.

## 2. Results

### 2.1. Genomic sequence-based language modelling captures emerging diversity in the SARS-CoV-2 spike gene

Our results validate the idea that a complex combination of codon weights may confer evolutionary advantage to the variant. The combinations of weights were learned using state-of-the-art unsupervised embeddings for capturing the latent space of spike DNA sequences of SARS-CoV-2. The framework of *Strainflow* is depicted in the figure below (*Figure 2A)*. The global tSNE plot represents dynamic emerging patterns derived from latent space representations of spike genes of SARS-CoV-2 (*Figure 2B*) from September, 2020 to March, 2021, along with specific geographic locations (country-level) such as India, UK, USA, and Brazil.

**Figure 2.**
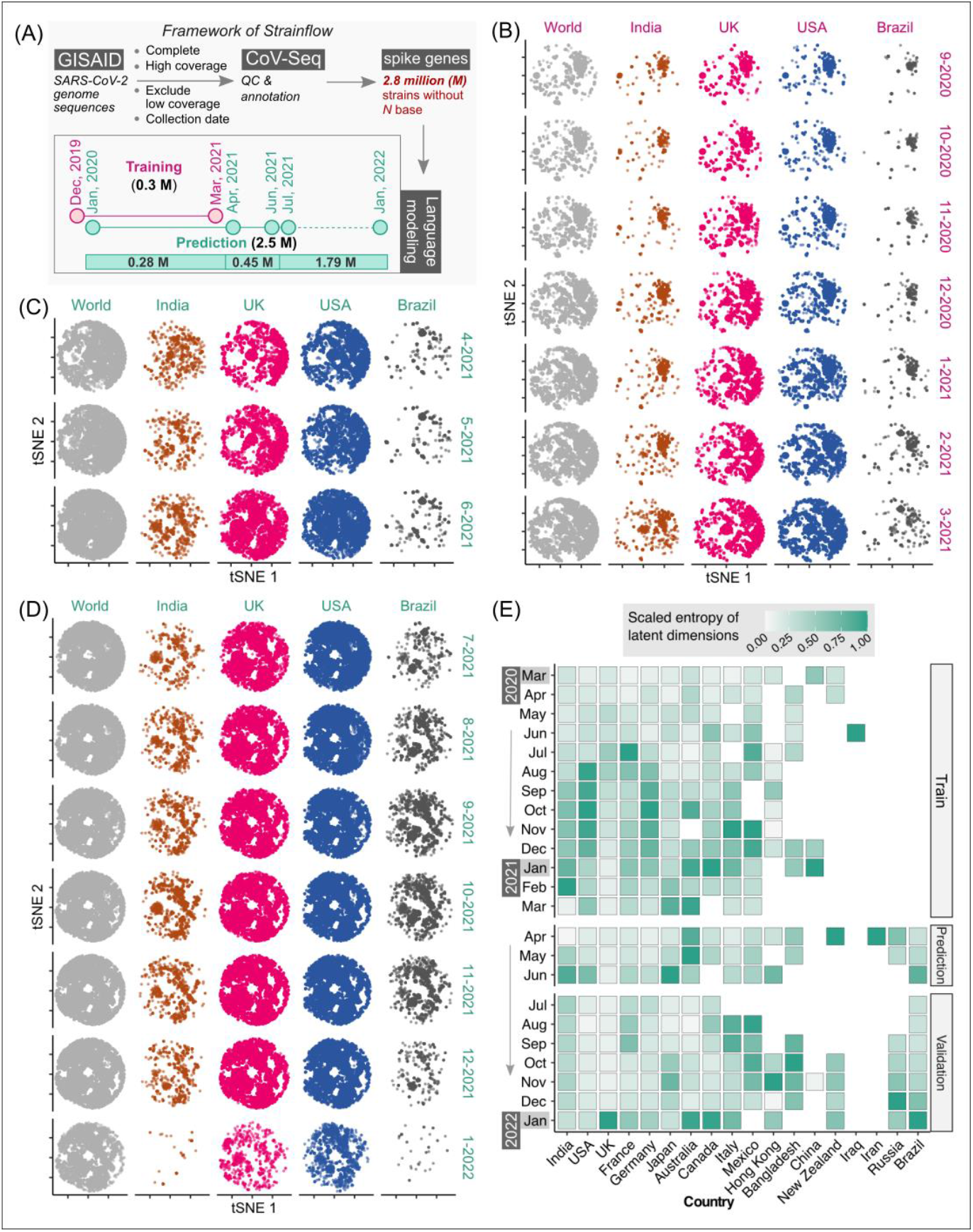
Latent space of spike genes derived using Strainflow preserves spatiotemporal information of SARS-CoV-2 spread. (A) The implementation framework of Strainflow (details described in the method section) (B) tSNE plot showing distinct spatio-temporal relationship based on the latent space learned from the spike gene of 0·308 million SARS-CoV-2 genomes collected till 31 March 2021 (world), India, UK, USA, and Brazil. (C) Embeddings estimated or predicted from the Strainflow model for 0·45 million SARS-CoV-2 spike genes from the month of April, 2021 to June, 2021. (D) Embeddings estimated or predicted from the Strainflow model for 1.79 million SARS-CoV-2 spike genes from the month of July, 2021 to January, 2022. (E) Heatmap showing the scaled entropy for 18 countries from March, 2020 to January, 2022 (showing data for a. training: March, 2020 to March, 2021, b. prediction: April, 2021 to June, 2021, and c. validation: July, 2021 to January, 2022). The entropies for each country were scaled to the same range to visualize the temporal trends within the country.

To investigate the information content in the latent space of the spike gene learned by our *Strainflow* pipeline, we performed qualitative and quantitative analysis on 2·7 million SARS-CoV-2 spike genes collected from December, 2019 to January, 2022. Qualitative analysis was performed by performing dimensionality reduction with a fast tSNE method called Flt-SNE (Linderman et al., 2019). We compared the 2D t-SNE plot of the world with four countries (India, UK, USA, Brazil) from September, 2020 to January, 2022, which clearly highlights the dynamic changes in the spike genes across countries in different months (*Figure 2B, 2C, 2D*). Additionally, quantitative analysis of the latent space was performed by calculating the fast sample entropy of each latent dimension (Tomčala, 2020). To compare the monthly entropy of the latent dimensions of different geographical regions, the mean entropy was calculated and normalized across the months for each country. We observed the highest entropy (information content) for India, UK, USA and Brazil in the months of February-2021, January-2022, August-2020, and January-2022 respectively. Interestingly, we observed high entropy for 4 months from August, 2020 to November, 2020 in the USA (*Figure 2E*). This highlights that the spike protein latent space representation learned by *Strainflow* could be used as a proxy to capture the spatiotemporal entropy or diversity in the emerging SARS-CoV-2 strains across different countries.

### 2.2. Preservation of spatiotemporal information of SARS-CoV-2 spread depicted with phylogenetic analysis

Sequence-level embeddings were obtained from the codon embeddings and investigated for the presence of genomically meaningful characteristics. The phylogenetic tree derived from the embeddings for the United Kingdom (*Figure 3A*) shows two clear temporally split clusters for 2020 and 2021 sequences, which may be indicative of different strains in these time periods. The temporality of the collected sequences was found to be preserved in the two clusters, although the model was trained only on genome sequences.

**Figure 3.**
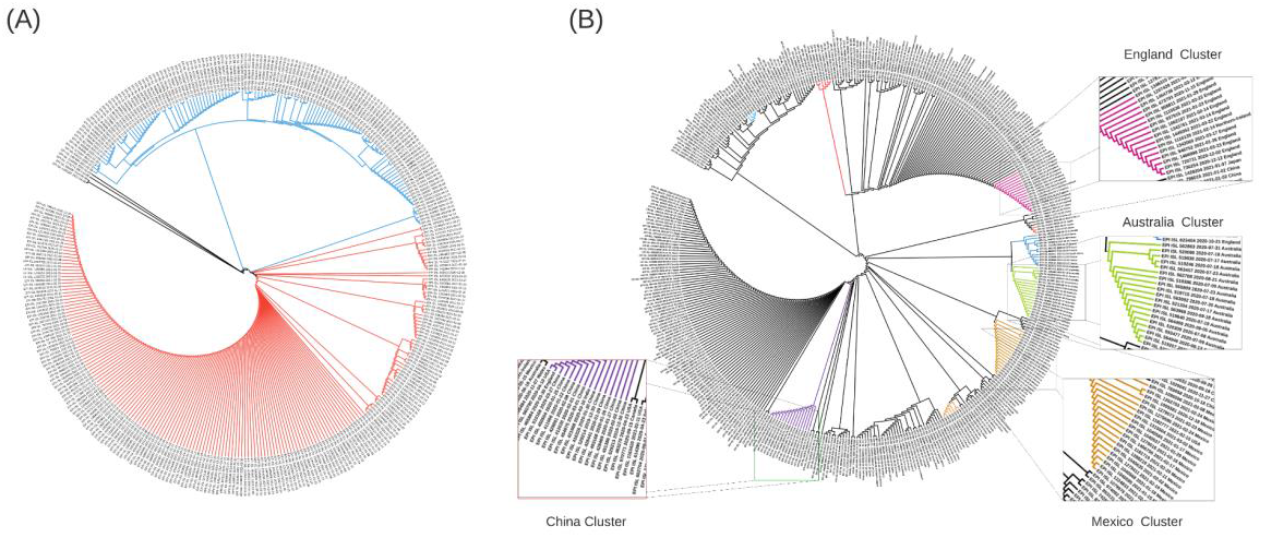
Phylogenetic trees constructed using cosine similarities between 400 randomly sampled sequence embeddings. (A) Dendrogram for strains from the U.K.: Cluster 1 (blue) contains strains from the period Oct 2020 - Dec 2020, while Cluster 2 (orange) contains strains collected between Jan 2021 - Mar 2021. (B) Dendrogram for 16 countries across the globe: Chinese, Australian and England strains form tight clusters (marked in purple, green, and magenta), while strains from Italy, France, Brazil, Japan, Canada, USA, Scotland, and India are dispersed with other countries.

The phylogenetic tree with globally collected sequences (*Figure 3B*) demonstrates that geospatial information is also preserved in the sequence embeddings. The dendrogram constructed using cosine distance between embeddings revealed clear clusters of geospatially close regions. Embeddings from geographically close locations were clustered together (Figure 3), and countries closer geographically had similar embedding patterns (Figure 4). This highlights that our de-novo embeddings captured these similarities without the need for standard alignment methods or expert knowledge of lineages. Clusters for China (purple), Australia (green), and England (magenta) are highlighted in Figure 3B. Strains from Italy, France, Brazil, Japan, Canada, USA, Scotland, and India were found to be dispersed with other countries. Overall, *Strainflow* captures the temporal emergence of strains and geographic information in a country-specific manner.

**Figure 4.**
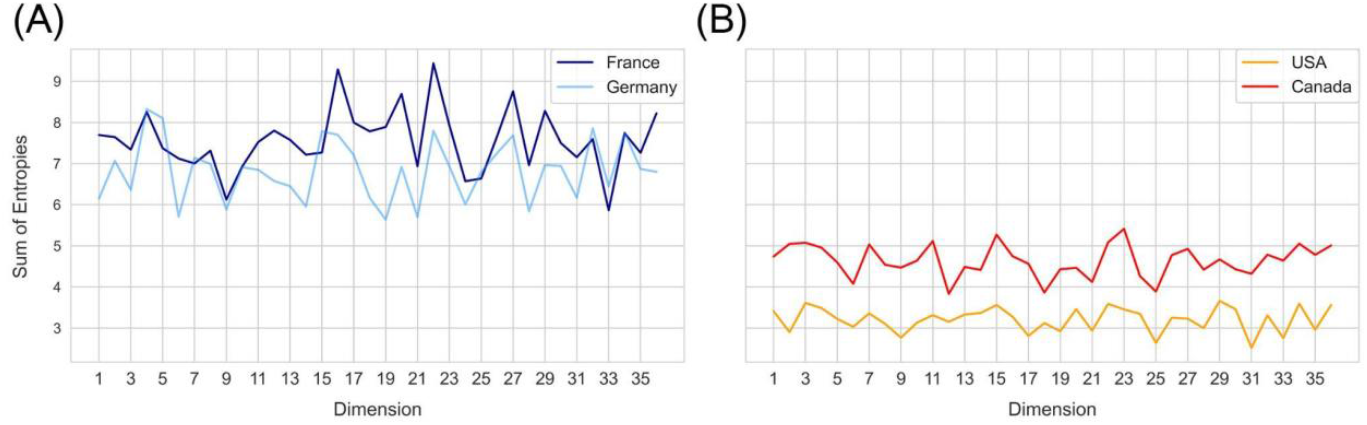
Sum of sample entropy for each latent dimension for different countries. Country pairs (**A**) - France and Germany, (**B**) USA and Canada show a similar distribution of total sample entropy across dimensions, while each pair differs from the other.

### 2.3. Entropy in the latent space dimensions captures variability in the spike gene

Entropy of a latent dimension has biological significance as it intuitively captures the variation in codon level changes during a certain time window. Each latent dimension encodes a combination of codon weights and increase in entropy represents frequent changes to these weights. Temporal changes in entropy are therefore expected to uncover the explore-exploit cycles of SARS-CoV-2 spike gene changes, hence biologically indicative of future trends. To compare different geographical regions, the sum of sample entropy was computed for each latent dimension across all the months. This revealed that certain geospatial regions such as France and Germany (*Figure 4A*) and USA and Canada (*Figure 4B*) have similar total entropies across the latent dimensions, indicating that strains in these regions have been accumulating similar genomic changes.

### 2.4. Entropy dimensions are predictive of new COVID-19 caseloads

We then attempted to decipher the relationship between monthly sample entropy and monthly new COVID-19 cases in different countries. Detrended cross-correlation coefficient was calculated at different lag values, which revealed that entropy dimensions have a leading relationship with new cases (*Figure 5A, 5B*). This suggests that the genome sequence data in a given month can be used to predict new cases in subsequent months. A lead period of two months was chosen and Boruta algorithm was employed to assign feature importance scores to different dimensions, which revealed that dimension 32 is the most significant predictor of new cases (*Figure 5C*). Significant dimensions from Boruta analysis were used for further modeling. Random forest based regression modelling on the predictive features achieved a total R-squared of 73% on the validation set. The predicted cases were found to be highly correlated with the actual cases (*Table 1*), which suggests that our model can indicate the directional change of cases for different countries. Further, the predicted relative change in cases between successive months was found to be correlated to the actual relative changes (*Table S1*), which suggests that our model can also indicate the magnitude of change that we expect to observe in the cases.

**Table 1.**
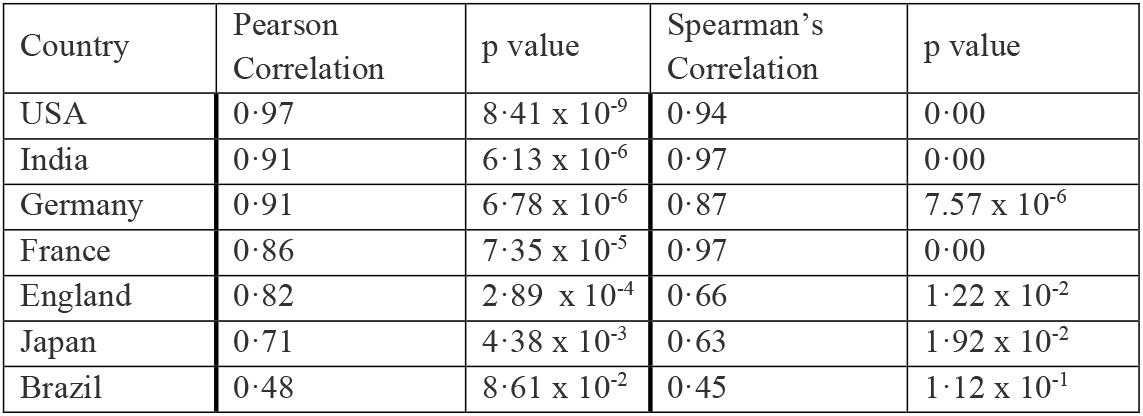
Pearson and Spearman’s Correlation coefficients between predicted and actual cases in different countries.

**Figure 5.**
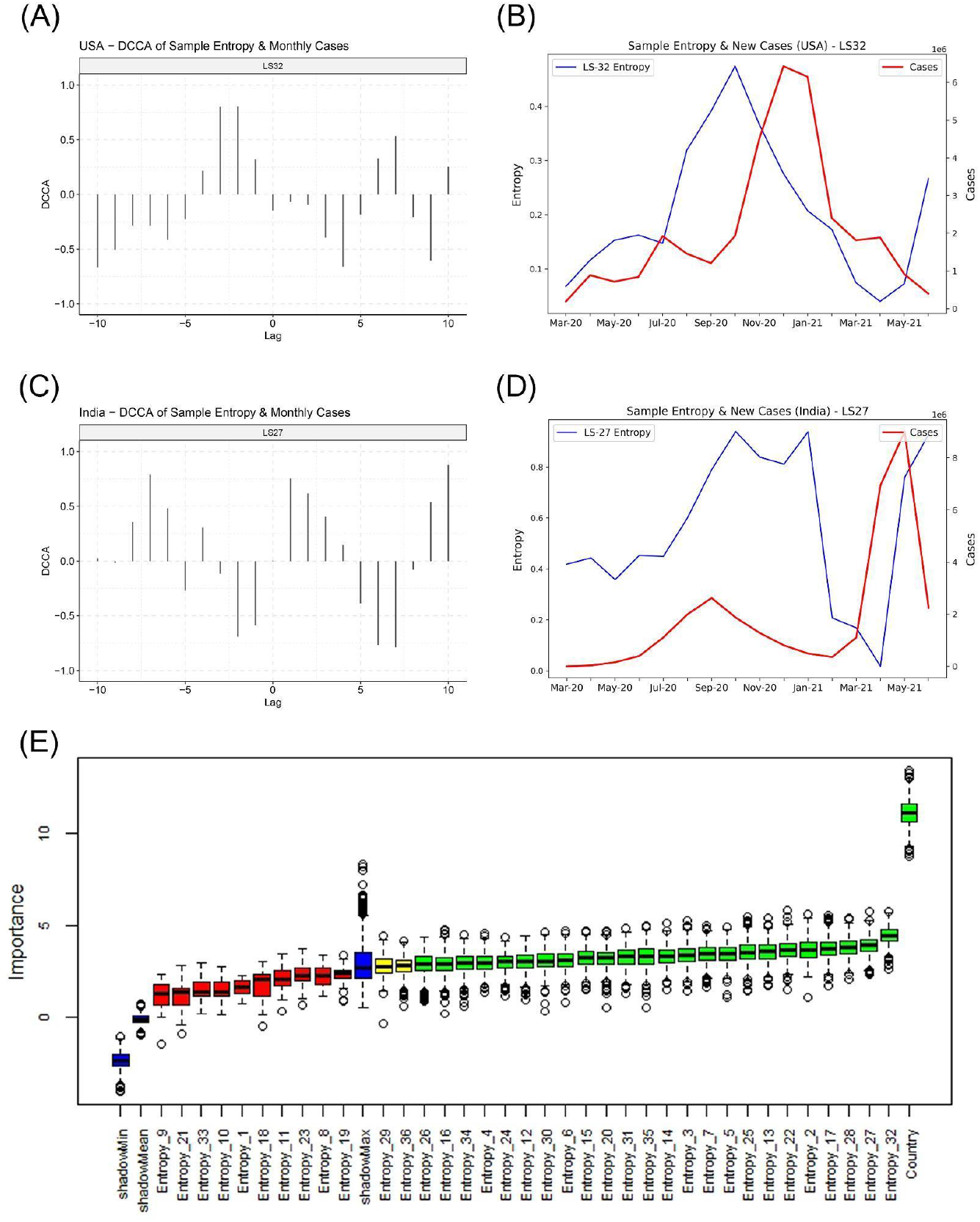
Relationship of the entropy of latent space dimensions with COVID-19 caseloads. (A) Detrended Cross-correlation coefficient values for different lags between Entropy dimension 32 and new cases for USA. High values are observed for a lead of 1 and 2 months. (B) Line plot for Sample Entropy dimension 32 and monthly new cases for USA, indicating that the entropy in dimension 32 has a leading relationship with the cases. (C) Detrended Cross-correlation coefficient values for different lags between Entropy dimension 27 and new cases for India. (D) Line plot for Sample Entropy dimension 27 and monthly new cases for India, indicating that the entropy in dimension 27 has a leading relationship with the cases. (E) Feature importance scores from the Boruta algorithm for predicting cases in the month following the next month.

Our model can be therefore used to predict the COVID-19 caseloads in several countries. Both USA (*Figure 6A*) and Japan (*Figure 6C*) show an increase in the sample entropy across the time period April - June 2021, concurrent with the respective spreads in these countries. Our model predicts new caseloads with a two-month lead time, which strongly predicts a spike in new cases both in USA (*Figure 6B*) and Japan (*Figure 6D*) in the months of July and August, 2021. For India our model predicted a decline in the number of cases for the month of July and August, 2021 (Figure 6E, 6F). Therefore our model may be used as an epidemiological early warning system to predict new caseloads.

**Figure 6.**
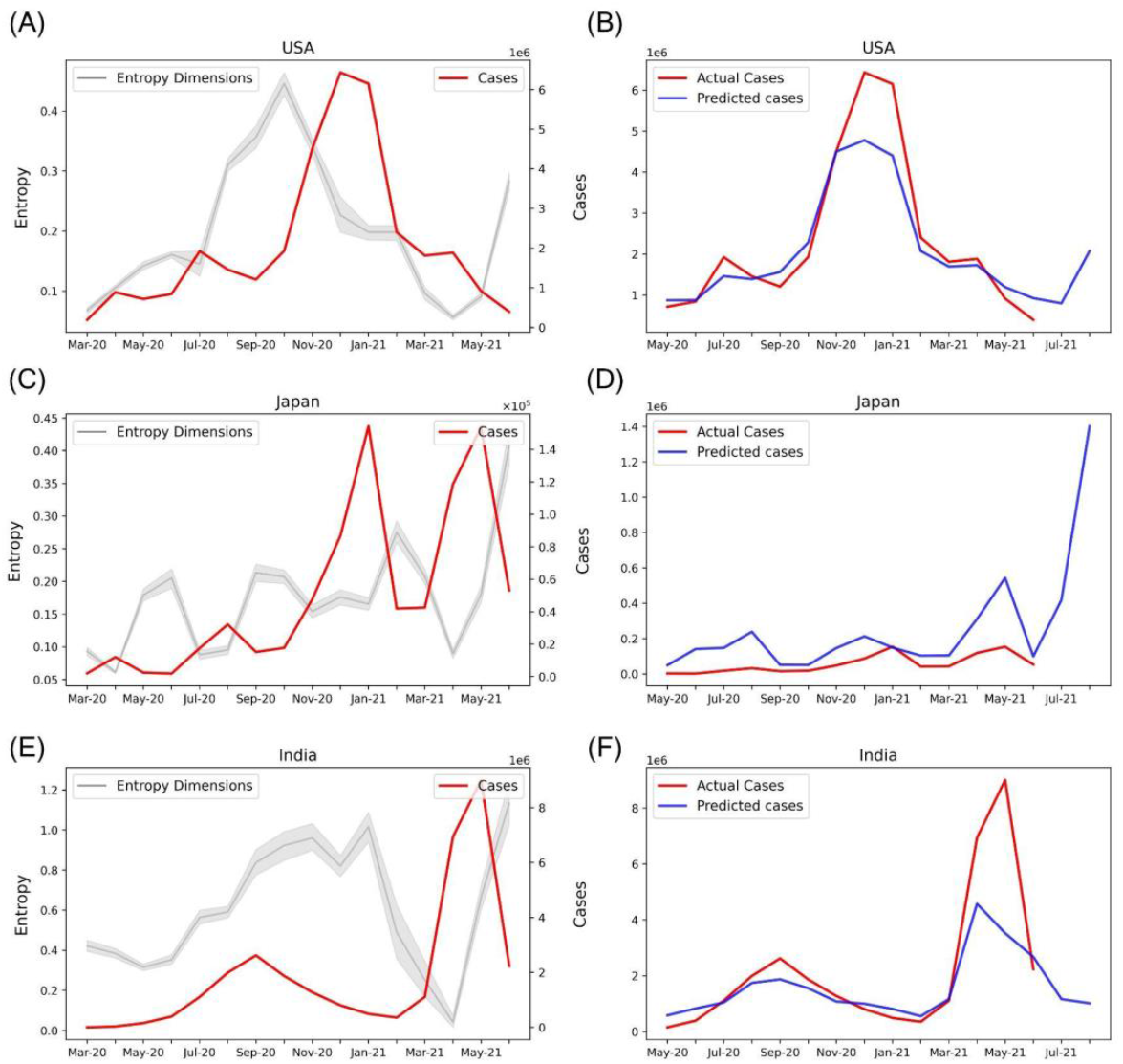
Prediction of new COVID-19 cases with Sample Entropy values of the latent dimensions. (A) Line plot showing the Entropy values of the selected features and new COVID-19 cases for the USA. (B) Actual and predicted cases based on the entropy values of selected features for the USA. The model predicts a rise in cases for July and August 2021. (C) Entropy of selected features and new cases for Japan. (D) Actual and predicted cases for Japan. A spike in cases is predicted for July and August 2021. (E) Entropy of selected features and new cases for India. (F) Actual and predicted cases for India.

### 2.5. Codons associated with the predictive features could be linked to SARS-CoV-2 variants

We further assessed the potential link of the predictive features with SARS-COV-2 variants by extracting the top 10 contributing codons and their associated weights for each dimension (*Table S2*). The intuition behind this idea is that the codons with high weights in a given dimension, when mutated in the viral sequence, are likely to cause a significant change in the entropy of the associated dimensions. Therefore each predictive feature can be linked to codons, which can further be mapped to Variants of concern (VOCs) and Variants of Interest (VOIs) (*Table S3*). Despite the fact that our model cannot directly capture the SARS-CoV-2 variants, it was observed that dimension-32 captures the CTG, CGG codons (ranks 5 and 8 respectively), known to be involved in the mutation T19R. Similarly, dimension 3 captures three codons (ACG, CGG, CAC) that are associated with multiple variants such as K417T, L452R, and D1118H, causing increased infectivity, pathogenicity, and spread. Dimension 30 captures codons CAT and CAC associated with Δ69 and D1118H respectively which are linked to B.1.1.7 lineage.

Codon weights (*Table S2*) of a given predictive feature provide an opportunity to associate with specific Variants of concern (VOCs) and Variants of Interest (VOIs), and to predict emerging SARS-CoV-2 variants. Distinct dimensions capture country-specific changes and may be surveilled to monitor the spread of the pandemic. This approach was back-validated with several real-world examples. For instance, dimension 32 captures the codons CGG (R) and CAC (R), which are found in B.1.429 lineage (L425R mutation). Dimension 3 captures CGG which is seen in L452R (associated with lineage B.1.617.1), which was first observed in India in December 2020 and was found to have increased infectivity and transmissibility.

### 2.6. Prospective validation of the model in the Delta and Omicron surges revealed interpretable predictive features

For investigating the potential of our predictive features to track the spread of SARS-CoV-2, we used the codon level information of the SARS-CoV-2 delta variant for the spike gene and extracted the weights of these codons specific to each feature. We selected Dimensions 3, 4, 12, 13, 15, 16, 25, 28, 30, 32 with high absolute weights for the codons related to the delta variants (*Figure 7A*). The entropy of these features was contrasted with the caseloads in England (*Figure 7B*), India (*Figure 7C*), and USA (*Figure 7D*). Overall, the temporal tracking of these features may be used as a surrogate to track the spread of various SAR-CoV-2 variants.

**Figure 7.**
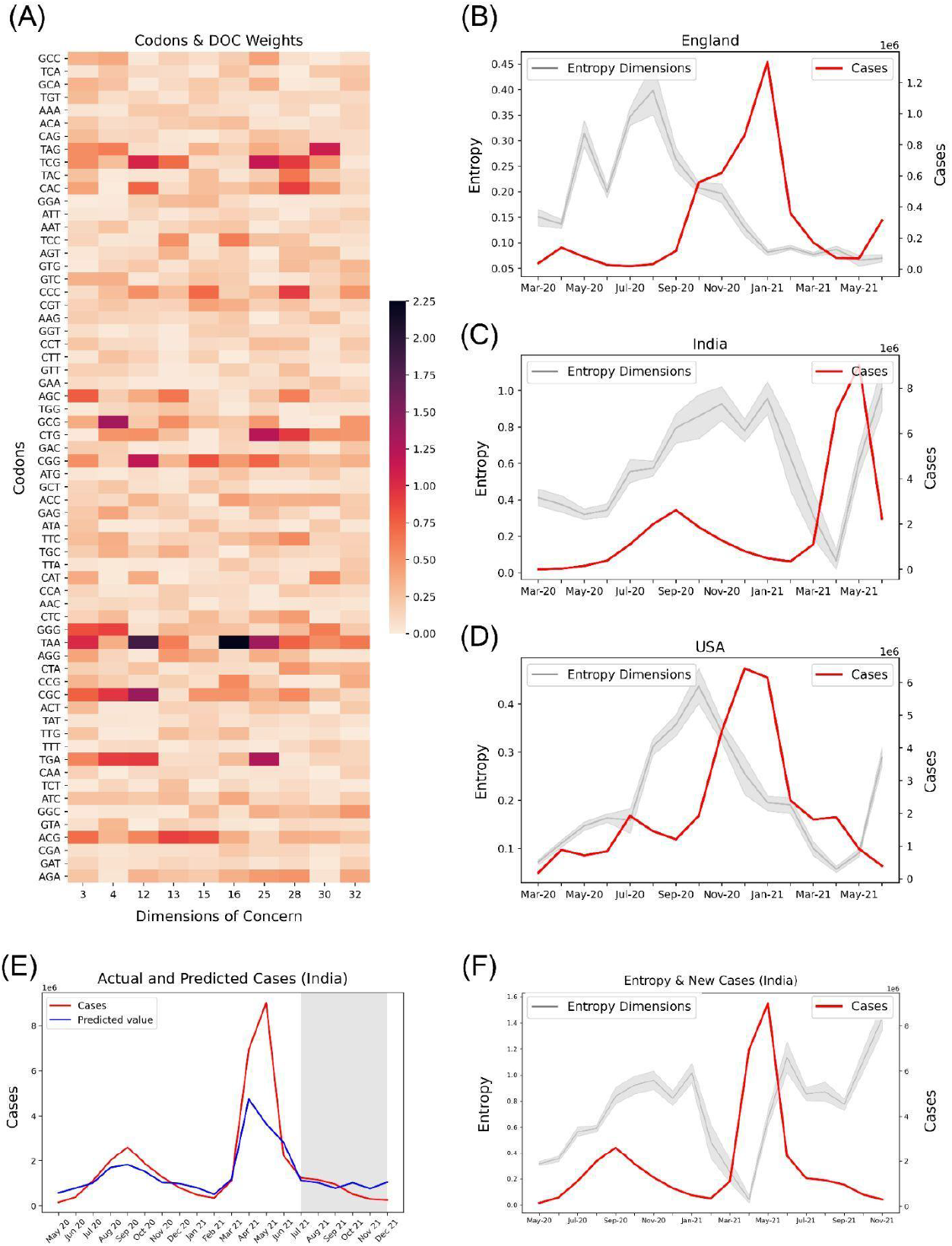
Potential association of codons observed in SARS-CoV-2 Delta variant (lineage B.1.617.2) with their corresponding entropy features, and the trend of caseloads with the entropy features. (**A**) Absolute latent space weights of the codons associated with the entropy features linked to the Delta variant. Line plots showing the entropy features and cases in countries, (**B**) England, (**C**) India, and (**D**) USA. The entropies show an increasing trend in the months April - June 2021 for India and USA, indicating a possible surge in the delta variant in these countries. (**E**) Predicted and Actual cases for India. The region shaded in grey represents the months for which the case prediction model was prospectively validated. (**F**) Entropy and Caseloads for India. Explore-exploit patterns in the genomic feature-space can be observed.

Our case prediction model was frozen in June, 2021 and prospectively predicted the caseloads from July, 2021 to December, 2021. Our model predicted the case upsurge in India due to the Omicron variant in November, and December, 2021 (*Figure 7E*) two months ahead of time. Although the model fails to predict the exact values of cases, it is useful as a trend indicator. Further, we observe explore-exploit cycles in the entropy-space of India prior to the case peak due to the Delta variant in May, 2021 (*Figure 7F*). A similar exploration phase can be observed for the months from September - November, 2021, which may be indicative of an upcoming case peak driven by the Omicron variant.

## 3. Discussion

We have implemented an approach for analyzing the emerging strains based on the latent space of spike protein coding nucleotide sequences. We chose the nucleotide sequences instead of proteins in order to capture and track the variations that may not have immediate functional consequences. Our approach has two main underlying tenets: (i) long-range interactions are known to modulate the functional interaction between receptor binding domain and ACE2 receptors, hence may be captured in the NLP models that capture 3-mer changes and context, and (ii) latent dimensions may be differentially correlated with indicators of spread, thus providing a data-driven handle for tracking and predicting variants of concern and variants of interest (Mugnai et al., 2020). The pipeline takes advantage of temporal changes in the semantics of mutating sequences. Preservation of phylogenetic structure based upon the similarity matrix obtained using the embeddings validated that the latent dimensions capture spatio-temporal information. Analyzing the dynamic patterns and underlying correlations in the 30,000 base pair long sequence of SARS-CoV-2 is important to highlight the mechanistic understanding of mutations (Shishir et al., 2021). SARS-CoV-2 seems to show a particularly high frequency of recombinations arising due to the absence of a proof-reading mechanism and sequence diversity, which calls for urgency in studying its transmission pattern (Rouchka et al., 2020; Mandal et al., 2021). Therefore predicting mutations in the spike protein, which binds to ACE2 receptors can help us estimate the spread of disease and the efficacy of therapeutic treatments and vaccines (Li et al., 2020; Srivastava et al., 2021).

While most research studies have attempted to predict the exact number of cases and have failed, our work is focussed on early prediction of trends from a non-obvious source of data. Unlike obvious data sources, the inter-relationships between codons in genome sequences are complex and less likely to be influenced or biased. Furthermore, sequencing data are made routinely available via various national and global consortia for genomic surveillance of SARS-CoV2. Our study also highlights the potential for triangulating insights from completely unrelated datasets, an approach that is expected to eliminate systematic biases in reporting by independent organizations. Further studies may triangulate insights from disparate, heterogeneous datasets such as mobility, genome surveillance, testing and case predictions to partially solve the problem of biases in individual datasets.

Entropy is a measure of the disorder of a system. We hypothesized that mutations increase the chaotic dynamics in the latent space of spike genes. To calculate entropy, we used the accelerated versions of the Approximate Entropy and Sample Entropy algorithms, called Fast Approximate Entropy and Fast Sample Entropy (Tomčala, 2020). Both algorithms aim to quantify how often different patterns of data are found in a time series. Fast Approximate Entropy, however, is a biased statistic and depends on the length of the series. Since we could have different counts of genome sequences collected each month, we preferred Sample Entropy, which is independent of the length of the series. Entropy values were calculated for each latent dimension in each month. Thereafter, Detrended Cross-Correlation Analysis (DCCA) was performed between the entropy dimensions and the new cases (Prass and Pumi, 2020b). DCCA is a modification of the standard cross-correlation analysis for finding relationships between non-stationary time series. High cross-correlation for different lead periods revealed that the entropy values in a given month could be used to predict the new cases in different countries in subsequent months. Different countries had different lead times at which the highest cross-correlation was observed between the entropy dimensions and the cases, ranging from 1-6 months. Overall, a lead time of two months was chosen to model the new cases. An empirical analysis was also done with daily values of entropy and new cases. Entropy was calculated in rolling windows, and cross-correlation analysis was performed between entropy and new cases at different lead periods. Although the cross-correlation values were found to be significant, the values were low and ranged between -0·1 to 0·1. Therefore, we decided to use the monthly entropy values for the modelling exercise.

To predict new COVID-19 cases, a random forest regression model was trained on the monthly entropy data. With sample entropy, we achieved an R-squared value of 73% on the validation set, while with approximate entropy, the value was only 10%. Therefore the model trained on sample entropy was selected. The predictions from the model were found to be highly correlated with the actual cases, indicating that our model can be used for preemptive warning signals for the rise in cases in different countries. Further, the actual and the predicted difference in the number of cases in consecutive months was found to be correlated, which suggests that the relative change in the cases in consecutive months predicted by our model is linked to the relative change in the number of cases. Overall, we recommend that our model be used to predict dangerous trends and not the actual number of cases. Further, the mapping from latent dimensions to Variants of Concern (VOCs) and Variants of Interest (VOIs) may help us track the country-specific spread of different variants.

The COVID-19 pandemic has been a dynamically evolving scenario, with new strains emerging and vaccines being developed. With the SARS-CoV-2 genome constantly mutating, we anticipate an underlying change in the grammar of the sequence, underpinning the need to update our language model every few months. Further, the regression model for caseloads needs to be periodically retrained too. An empirical analysis led us to discover that the Random forest model used for prospective validation from November, 2021 onwards performed better in terms of predicting the number of cases than the model used for prospective validation from July 2021 (*Figure S2*). However, both models indicate similar trends in cases for most countries.

Further, models trained on genomic sequences can be used for predicting infection severity based on Co-associations between the SNPs of Co-morbid Diseases and COVID-19 (Wang et al., 2020b). The machine learning models can also be trained on genomic sequences for COVID-19 classification (Arslan, 2021). Although the variance explained by our model is low, however, we were able to compute the variability associated with spike protein mutations. So our method showed a potential way to estimate the new cases variability associated with spike protein mutations. Our methods can be incorporated with the epidemic projections model to better predict the epidemic trajectories. The latent dimensions may further be employed to predict the clinical consequences of emerging strains. The currently available vaccines are intended for early SARS-CoV-2 strains, but with new emerging variants, immune responses triggered by these vaccines may be weaker and short-lived. As seen in the devastating second wave of the pandemic in India, newer SARS-CoV-2 variants have acquired an increased pathogenic potential resulting in rapid clinical progression and overwhelmed health systems. Mitigating such events in the future will require stronger surveillance systems in place. Our study offers a promising solution in this direction and lays the foundation for proactive genomic surveillance of COVID-19.

Our study has the following limitations. Our approach of codon embeddings does not indicate the position where the codon change may have happened in the spike gene. This is because low-dimensional embeddings do not preserve the positional encoding of words. However, we are investigating advanced approaches such as complex-valued word embeddings with positional encodings and transformer models such as BERT to overcome our current limitations (Lee et al., 2020; Wang et al., 2020a; Wolf et al., 2020). The latter are considered expensive and data-hungry models and it will remain to be evaluated if the gain of positional information may be countered by the loss of prediction accuracy for forecasting new cases in the future. However, we believe that the availability of sequences for a wide variety of viral pathogens presents an exciting opportunity to train data-hungry models that may be able to transfer insights across pathogens and yet remain interpretable. Further, our Strainflow model is trained only on the spike gene of the viral genome, which does not represent the complete variation spectrum of the virus. To mitigate this shortcoming, we will develop a genome-level *Strainflow* pipeline for SARS-CoV-2. Furthermore, the present study does not consider the interaction between the spike gene and other genes in the SARS-CoV-2 genome. We have not considered the interaction between the ACE2 receptor sequence for the human and the spike gene sequences due to the unavailability of such large-scale paired data. However, we believe this is a strength of our study as we were able to extract relevant features as well as make valid predictions using the spike region of the SARS-CoV-2 gene alone.

Our current approach does not explicitly capture specific positional mutations. Although the ad-hoc analysis for codon weights on significant dimensions allows us to rank the codon level changes, the predictive feature is a complex nonlinear combination of these changes which may eliminate strongly associated features. The E484Q mutation was not captured as the most important in our model. However, this may be because other codon level changes such as L452R and their combinations may be correlated and hence a proxy for E484Q. Importantly, the B.1.617 variant has both L452R and E484Q mutations and L452R change was predictive and captured in the top ten ranks for multiple latent dimensions (3, 4, 10, 12, 13, 15, 16).

Finally, a relatively small number of samples were used to construct the supervised predictive model for case prediction. As more data becomes available in subsequent months, we can produce more confident case predictions. An empirical validation depicted that we require a minimum of 100 samples per month for calculating the sample entropy. This also underscores the need for a more reliable and agile approach to deposit country-level datasets on repositories such as GISAID. We make an appeal to the countries to facilitate the sharing of such data in order to be prepared for any future waves of the current pandemic and for preventing the new emergence of strains. We believe our study is an instance of the new paradigm of pathogen surveillance using a novel language modelling approach that is potentially scalable to infectious disease surveillance and antimicrobial resistance.

## 4. Methods

### 4.1. Datasets

#### Training dataset

The dataset was downloaded from GISAID EpiCoV (April 8, 2021 release) (Shu and McCauley, 2017). 0·36 million genome sequences (December, 2019 - June, 2021) with high nucleotide completeness, coverage, complete temporal information, and presence of less than 5% non-identified nucleotide bases (N) were downloaded. The sequences included 63 countries, including India, United Kingdom, USA, Australia, New Zealand, Germany, Russia, Italy, France, Mexico, Canada, China, Japan, Pakistan, Bangladesh, Iran, Iraq, the continent of South America, and Africa. Duplicate samples were removed, and whole genome sequences were parsed using CoV-Seq to extract nucleotide sequences corresponding to each of the 12 Coding DNA Sequences (Liu et al., 2020). Accession IDs that did not cover 12 coding regions were discarded, yielding 0·31 million high-quality SARS-CoV-2 genome sequences for language modelling. The spike gene region of each sequence was filtered and used for all subsequent analysis. We downloaded country-wise COVID-19 data for new cases from a publicly available repository maintained by Johns Hopkins University Center for Systems Science and Engineering (JHU CSSE).

#### Evaluation dataset

We downloaded around 0·6 million genome sequences submitted to GISAID from April 2021 to June 2021. We used our trained model to predict the latent representations for these sequences.

### 4.2. Word Embeddings in Strainflow pipeline

In our Strainflow pipeline, we have adopted a word2vec model (Mikolov et al., 2013). Low dimensional representations for the genome sequences were learned using the word2vec model. Non-overlapping sequences of 3-mers (codons) were considered as words for training the model, which was implemented in Gensim (Řehůřek and Sojka, 2010). The skip-gram algorithm was used, with a fixed window size of twenty and vector size of thirty-six. For generating a consensus embedding for a particular strain, genomic sequences were represented by taking the mean of each codon occurring in the sequence dimension-wise. The mean was calculated by summing across all the k-mers over each dimension and then dividing it by the total number of codons present in the sequence. For selecting the dimension size for our word embeddings, we calculated the PIP (Pairwise Inner Product) loss (Yin and Shen, 2018). PIP loss is a metric used for calculating the dissimilarity between two word embedding matrices. For the embedding matrix of strains (E), the PIP matrix is defined as the dot product of the embedding matrix with its transpose (*E. E* ^*T*^). The PIP loss between two embedding matrices is defined as the norm of the difference between their PIP matrices.

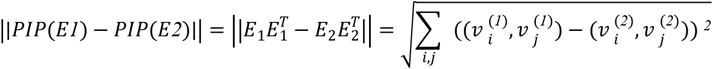

Various word2vec models were trained on the dataset with different vector sizes varying in multiples of three. Based on the PIP loss calculations, we found out that word embeddings with 36 dimensions showed a differential dent in the curve (change in straight line), due to which we selected this to be the dimension of the word embeddings (*Figure S1*).

### 4.3. Phylogenetic analysis using the latent dimensions of the spike genes

To evaluate the phylogenetic properties based on the latent dimensions of the spike gene, we computed the cosine distances among spike genes of SARS-CoV-2 with the 36 latent dimensions. The pairwise distance was further used for hierarchical clustering using the ‘hclust’ function in R statistical programming language. This analysis was performed using 400 random sequences of spike genes from 16 countries. The visualization of the phylogenetic tree derived based on the latent dimensions was done using ‘iTOL’ software (*Figure 2*) (Letunic and Bork, 2021).

### 4.4. Entropy of the latent dimensions

To quantify the properties of latent dimensions, we have used a well-known information theory based algorithm suitable for time series datasets, called ‘Fast Sample Entropy’(Pan et al., 2011). To compute Fast Sample Entropy, we have used the ‘FastSampEn’ function in the ‘TSEntropies’ package in R (Tomcala, 2018). Fast Sample Entropy can be computed as follows.

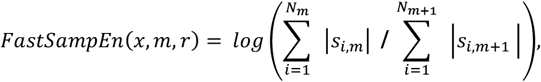

where,

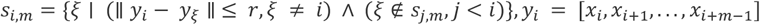

*s*_*i,m*_ is a set of sub-sequences of length m belonging to the i-th neighbourhood, and *N*_*m*_ is the number of these neighbourhoods.

In our case, ‘x’ is the latent dimension of the spike genes of the SARS-CoV-2 strains per month for a given country with default values of ‘m’ and ‘r’. Entropy was computed for each latent dimension on a monthly basis for each country. To compare geographies across months, we used average entropy derived from 36 latent dimensions, followed by normalization using all the monthly entropies for a given country (*Figure 1D*). To compare the entropy of the latent dimensions among countries, we used the total entropy of the country for each dimension and visualized it with line graphs (*Figure 3*).

### 4.5. Detrended Cross Correlations Analysis (DCCA)

To investigate the information content (entropy) of the latent dimensions with the new cases observed for COVID-19, we used the Detrended Cross Correlation Analysis(Prass and Pumi, 2020b) Here, DCCA captures the long-range cross correlation between time series (entropy of the months and caseloads for a given country). We tested both time series for stationarity using Augmented Dickey-Fuller (ADF) test (Mushtaq, 2011). The ADF test was implemented using the function ‘adf.test’ available in the ‘tseries’ package in R (Trapletti et al., 2020). Due to the non-stationary distribution of the estimated entropies and the caseloads for a given country, we used the ‘DCCA’ R package (Prass and Pumi, 2020a). Cross-correlation was calculated between the entropy dimensions at time t+h and new cases at time t, where h = 0, ±1, ±2, ±3 … ±10.

### 4.6. Machine Learning based identification of significant predictive features

Country-wise monthly total new cases data was taken at the end of each month. Total new cases data for each month was merged to monthly entropy dimensions data from March, 2020 to June, 2021. We used a regression based machine learning approach called ‘Boruta’, a wrapper algorithm around a random forest algorithm to select the most relevant entropy dimensions for the prediction of subsequent two months’ new cases(Kursa et al., 2010; Kursa and Rudnicki, 2020). We used the default parameters with the modification of the maximum runs as 1000. We selected the confirmed entropy dimensions as the most relevant predictive features for the prediction of new-cases.

### 4.7. Model development and evaluation for prediction of new cases in subsequent months

To predict the new cases in the next to next months, we used a regression based random forest model using the most relevant predictive features using the ‘Boruta’ R package (Kursa and Rudnicki, 2020). The model training was performed using entropy data from March, 2020 to February, 2021; and the fitted model was validated on entropy data from March, 2021 to April, 2021. The regression modelling was performed using 1000 decision trees using the ‘randomForest’ package in R (Liaw and Wiener, 2002).

### 4.8. Top codons associated with predictive features

To find the top codons associated with the latent dimensions, we extracted the absolute weights of each codon for a given dimension. The top 10 codons having the highest absolute weights (contribution) were identified corresponding to each dimension to link these to SARS-CoV-2 variants. We collected the SARS-CoV-2 variants and their associated genetic variations at the codon level linked to the spike gene, and a list of codons associated with VOIs and VOCs was curated (Table S4)(Lopez-Rincon et al., 2021; Naveca et al., 2021; Peacock et al., 2021; Srivastava et al., 2021; CDC, 2022). The curated list is based on the CDC guidelines, and we are consistent with their definition of lineage and variant (CDC, 2022).

### 4.9. Strainflow algorithm

The algorithm for the Strainflow pipeline has been described below:

1. We have collected the SARS-CoV-2 sequences from the GISAID EpiCoV database. High quality sequences with complete temporal information were filtered.
2. We extracted the spike gene region of these sequences from FASTA files using the CoV-Seq tool. A CSV containing these sequences and other metadata such as country names and dates was created.
3. The sequences were splitted into chunks of three characters (codons). A splitted sequence represents a document with three-letter words.
4. We trained a word2Vec model on the spike gene sequences for learning 36-dimensional word embeddings. The average of all word embeddings in a given sequence was treated as the embedding of the sequence.
5. We calculated the sample entropies of each dimension of our embeddings for each month and country.
6. New COVID-19 cases for each country in each month were calculated using data from the JHU CSSE repository.
7. A feature selection algorithm (Boruta) was used for selecting the entropy dimensions predictive of caseloads two months in advance.
8. Random Forest regression algorithm was used for predicting new cases two months ahead of time. The inputs to the model are the country names and important features extracted from the Boruta algorithm. The predictor variable is the caseload two months ahead of time for each country.

## Acknowledgements

This work was supported by the Delhi Cluster-Delhi Research Implementation and Innovation (DRIIV) Project supported by the Principal Scientific Advisor Office, Prn.SA/Delhi/ Hub/2018(C) and the Center of Excellence in Healthcare supported by Delhi Knowledge Development Foundation (DKDF) at IIIT-Delhi. We also thank Dr. Chitra Pattabiraman (NIMHANS) for her valuable inputs on viral sequences, and Harleen Kaur, Nishkarsh Saxena, and Anjali for their contribution to the dashboard visualizations.

## Data and Code Availability Statement

The SARS-CoV-2 sequence dataset was downloaded from the GISAID database (https://www.epicov.org/epi3/start). This study does not involve any human participants and all the data that have been used is in accordance with terms and compliance from the GISAID community. Based on the terms and conditions provided by GISAID, we cannot share the sequence dataset. However, the accession IDs of the SARS-CoV-2 strains are provided along with this manuscript. The latent representation dataset is available on request from the Lead Contacts. The code related to this study is available at: https://github.com/rintukutum/strainflow_manuscript

## Author contributions

Conceptualization: S.N., R.P., S.T., S.A., T.S.

Methodology: S.N., R.P., A, A.T., A.N., S.T., R.K., T.S.

Investigation: S.N., R.P., A, A.T., A.N., S.T., R.K., H.P.

Visualization: S.N., R.P., R.K., A, A.T., A.N., S.A.

Dashboard creation: R.M.

Project administration: T.S., R.K.

Supervision: T.S., R.K.

Writing - original draft: S.N., R.P., R.K., S.T., A, A.T., A.N., H.P.

Writing - review & editing: S.N., R.P., S.A, S.T, H.P., R.K., T.S.

## Strainflow Dashboard

*Strainflow*, our web application with results of online tracking and prediction of COVID-19 caseloads is available publicly at http://strainflow.tavlab.iiitd.edu.in/. The predictive models will be retrained with the new SARS-CoV-2 genomic sequences for the upcoming months.

