## Supplementary for "Genomic Surveillance of COVID-19 Variants with Language Models and Machine Learning"

### Supplementary Information

#### *Supplementary text*

#### Entropy as a measure of the information content of the latent dimensions

We hypothesized that mutations increase the chaotic dynamics in the latent space of spike genes. To calculate entropy, we used the accelerated versions of the Approximate Entropy and Sample Entropy algorithms, called Fast Approximate Entropy and Fast Sample Entropy (Tomčala, 2020). Both algorithms aim to quantify how often different patterns of data are found in a time series. Fast Approximate Entropy, however, is a biased statistic and depends on the length of the series. Since we could have different counts of genome sequences collected each month, we preferred Sample Entropy, which is independent of the length of the series.

#### Monthly entropy and DCCA to model the new cases

Detrended Cross-Correlation Analysis (DCCA) was performed between the entropy dimensions and the new cases (Prass & Pumi, 2020b). DCCA is a modification of the standard cross-correlation analysis for finding relationships between non-stationary time series. High cross-correlation for different lead periods revealed that the entropy values in a given month could be used to predict the new cases in different countries in subsequent months. Different countries had different lead times at which the highest cross-correlation was observed between the entropy dimensions and the cases, ranging from 1-6 months. Overall, a lead time of two months was chosen to model the new cases.

A similar analysis was done with daily values of entropy and new cases. Entropy was calculated in rolling windows, and cross-correlation analysis was performed between entropy and new cases at different lead periods. Although the cross-correlation values were found to be significant, the values were low and ranged between -0·1 to 0·1. Therefore, we decided to use the monthly entropy values for the modeling exercise.

#### Prediction of new cases with ‘blips’

We also experimented with “blips” as a feature to predict the new cases. Blips are sudden changes in the values of the latent dimensions. These changes may be caused by a mutation, which changes the words (codons) in a given genome sequence. This hypothesis was validated in simulation experiments in synthetic datasets. Each dimension of the spike gene embeddings for a country was analyzed for the presence of temporal anomalies. Countries having a minimum of 20 samples in any given month were selected and the same number of records (minimum samples in any given month for that country) were sampled without replacement from each month. These records were used to define control limits of ±1 standard deviation from the mean value for each dimension, and all values in the full dataset outside those limits were categorized as ‘Blip’ points. Blip counts in each month were normalized by calculating the number of blips per sample collected in a month for each dimension (normalized blips). The embedding dimensions were then compared in terms of the total normalized blips for each country to observe the significant dimensions and dis(similarity) in trends among different countries. Cumulative counts of normalized blips were analyzed to understand the temporal accumulation of blips in each dimension. Similar to entropy, blips were found to have a leading relationship with the cases. However, regression modelling results with sample entropy were found to be better.

### Strainflow Dashboard

*Implementation:* The strainflow dashboard web application is primarily built using *ReactJS* and other accompanying libraries for UI needs and *GraphJS* for graphical needs. Python libraries such as numpy, pandas, matplotlib and seaborn were used to pre-process and infer the dataset. The Random Forest regression model was implemented using the R library *randomForest*. The web application is available for use on http://strainflow.tavlab.iiitd.edu.in and works on all modern browsers.

*Functionalities:* The application has three tabs: Cases Plots, Entropy Plots, and Paper. The Cases Plot tab exhibits two graphs; one compares the actual number of cases with our predicted cases, with a two-month lead time, while the second shows entropy against the caseload for a given country.  The Entropy Plots tab displays the sum of sample entropy across all the latent dimensions for each pair of countries. The toggler present above the graph can be used to change countries to compare their entropies. Lastly, the “Paper” tab presents a graphical abstraction of our paper.

*Discussion*: COVID-19 had a devastating impact on our health systems, thus with caseload predictions made two months in advance, we provide a data-driven handle on epidemiological surveillance to warn about potential upcoming case surges, so that people can be prepared in advance and appropriate preemptive steps can be taken by policymakers to prevent the spread of COVID-19.

**Supplementary Figures**

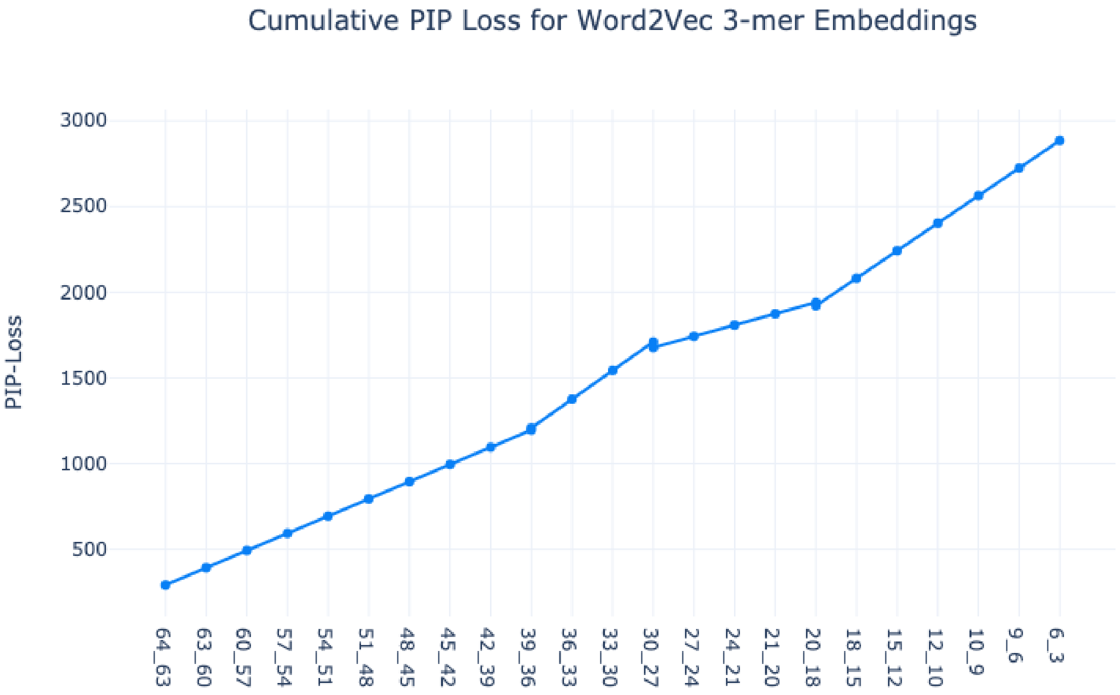


**Figure S1.** ***Determination of optimal dimensions in Word2Vec model****: Cumulative PIP Loss computed between word2vec codon embeddings differed by vector size taken in multiples of 3, till dimension size 64.*

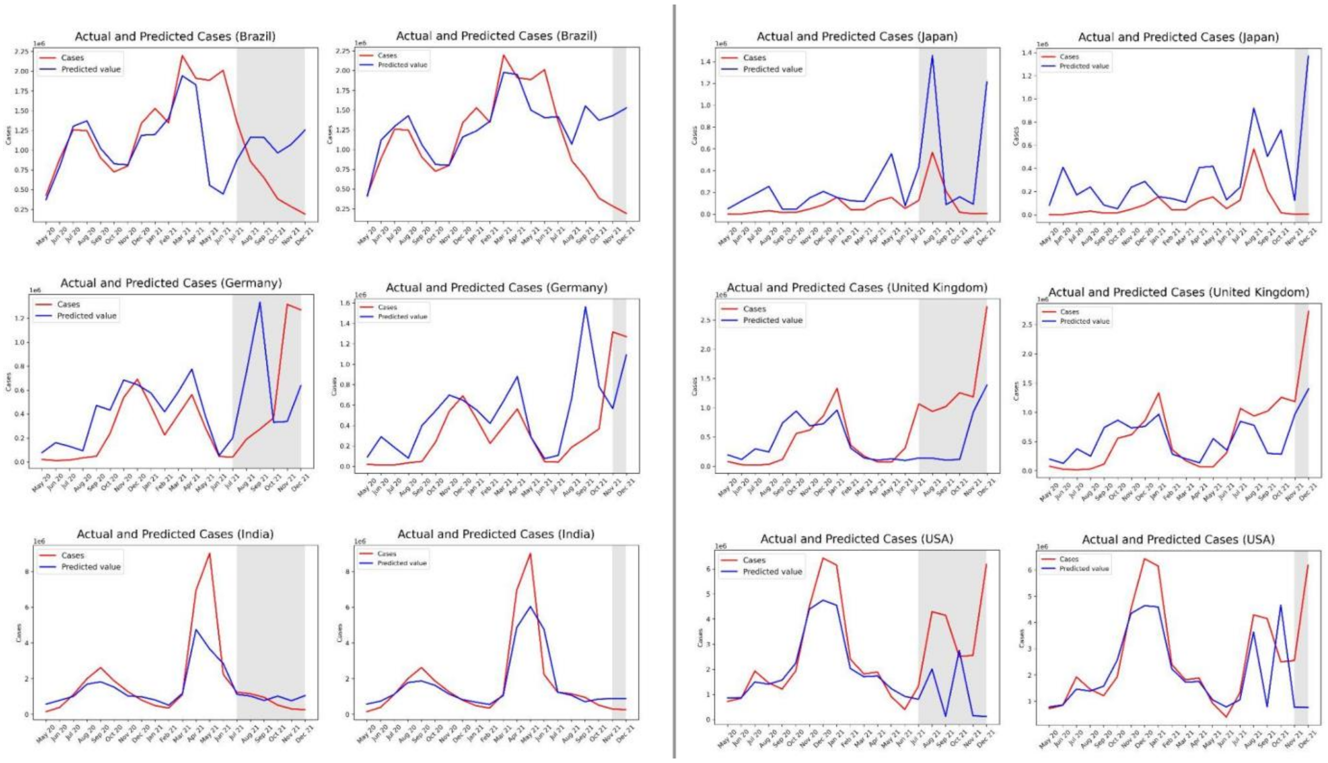

**Figure S2.** ***Comparison of regression model frozen in June 2021 compared to the one frozen in October 2021 for all countries.***

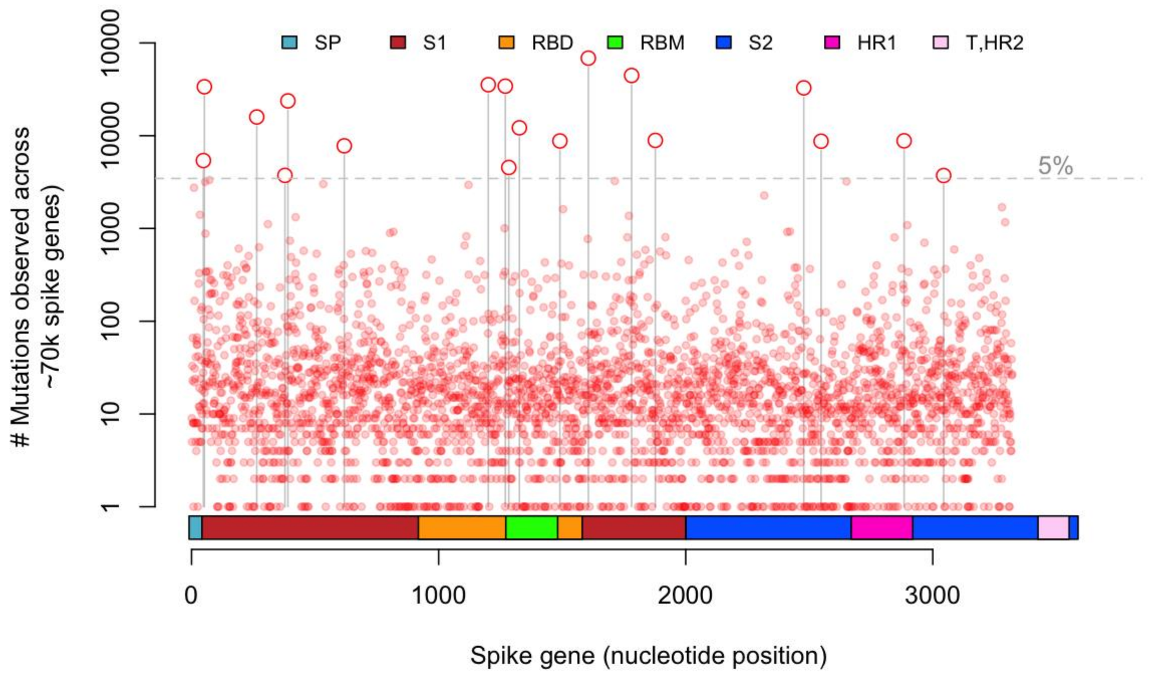


**Figure S3.** ***Spike gene domain structure and associated mutations till Jan, 2022.***

### Supplementary Tables

| Country | Pearson Correlation | p value | Spearman's Correlation | p value |
|---|---|---|---|---|
| USA | 0.93 | $2.81 \times 10^{-6}$ | 0.94 | 0.00 |
| England | 0.82 | $5.60 \times 10^{-4}$ | 0.71 | $8.14 \times 10^{-3}$ |
| France | 0.8 | $9.36 \times 10^{-4}$ | 0.55 | $5.25 \times 10^{-2}$ |
| Germany | 0.76 | $2.47 \times 10^{-3}$ | 0.77 | $2.92 \times 10^{-3}$ |
| India | 0.71 | $7.05 \times 10^{-3}$ | 0.59 | $3.60 \times 10^{-2}$ |
| Japan | 0.68 | $1.08 \times 10^{-2}$ | 0.57 | $4.73 \times 10^{-2}$ |
| Brazil | 0.67 | $1.22 \times 10^{-2}$ | 0.78 | $2.62 \times 10^{-3}$ |

**Table S1**. ***Pearson and Spearman Correlation coefficients between the delta of predicted and actual cases in different countries.***

Latent Dimensions from Word Embeddings

| 32 | | 28 | | 13 | | 3 | | 15 | | 30 | |
|---|---|---|---|---|---|---|---|---|---|---|---|
| **3-mers** | **Weights** | **3-mers** | **Weights** | **3-mers** | **Weights** | **3-mers** | **Weights** | **3-mers** | **Weights** | **3-mers** | **Weights** |

| | | | | | | | | | | | |
|---|---|---|---|---|---|---|---|---|---|---|---|
| TAA | 0.64 | CTG | 0.93 | ACG | 0.9 | TAA | 1.02 | ACG | 0.84 | TAG | 1.11 |
| CCC | 0.49 | TCG | 0.93 | TCG | 0.7 | GGG | 0.83 | CGG | 0.81 | GGG | 0.62 |
| GCG | 0.49 | CCC | 0.92 | AGC | 0.65 | AGC | 0.77 | CCC | 0.71 | CAT | 0.55 |
| GGC | 0.48 | CAC | 0.91 | TAA | 0.61 | CGC | 0.75 | CGC | 0.5 | TAA | 0.53 |
| CTG | 0.48 | TAA | 0.74 | GCG | 0.53 | ACG | 0.69 | CGT | 0.42 | CTG | 0.5 |
| AGA | 0.4 | TAC | 0.65 | TCC | 0.51 | TGA | 0.58 | AGG | 0.35 | TCG | 0.46 |
| CCG | 0.37 | AGC | 0.6 | AGG | 0.47 | TAG | 0.53 | TAG | 0.33 | CAC | 0.45 |
| CGG | 0.34 | TTC | 0.55 | TGC | 0.35 | TCG | 0.53 | GGA | 0.3 | ACC | 0.35 |
| CTA | 0.33 | CGC | 0.55 | AGT | 0.32 | CGG | 0.52 | CTC | 0.29 | ACG | 0.33 |
| GTG | 0.3 | AGA | 0.5 | CGG | 0.32 | CAC | 0.42 | GGC | 0.28 | CTA | 0.3 |

**Table S2: Dimensions of Concern and top 10 codons with their absolute weights.**

| DOC | Rank | Codon | Amino Acid | Variant |
|---|---|---|---|---|
| 32 | 5 | CTG | (Leu/L) Leucine | L452R |
| | 6 | AGA | (Arg/R) Arginine | T19R, R158G |
| | 8 | CGG | (Arg/R) Arginine | L452R |
| 27 | 1 | AGC | (Ser/S) Serine | R190S |
| | 7 | CTG | (Leu/L) Leucine | L452R |
| | 10 | CGG | (Arg/R) Arginine | L452R |
| 28 | 1 | CTG | (Leu/L) Leucine | L452R |
| | 4 | CAC | (His/H) Histidine | Q1071H, D1118H, Q677H |
| | 7 | AGC | (Ser/S) Serine | R190S |
| | 8 | TTC | (Phe/F) Phenylalanine | V367F, Del157 |
| | 10 | AGA | (Arg/R) Arginine | T19R, R158G |
| 17 | 1 | CGG | (Arg/R) Arginine | L452R |
| | 2 | CTG | (Leu/L) Leucine | L452R |
| | 6 | ACG | (Thr/T) Threonine | K417T |
| | 10 | GCC | (Ala/A) Alanine | A475H |
| 2 | 2 | CGG | (Arg/R) Arginine | L452R |
| | 3 | CTG | (Leu/L) Leucine | L452R |
| | 9 | CAC | (His/H) Histidine | Q1071H, D1118H, Q677H |
| 22 | 3 | ACG | (Thr/T) Threonine | K417T |

| | | | | |
|---|---|---|---|---|
| | 8 | AGT | (Ser/S) Serine | R190S |
| | 9 | CGG | (Arg/R) Arginine | L452R |
| 13 | 1 | ACG | (Thr/T) Threonine | K417T |
| | 7 | AGG | (Arg/R) Arginine | R190S |
| | 9 | AGT | (Ser/S) Serine | R190S |
| | 10 | CGG | (Arg/R) Arginine | L452R |
| 25 | 3 | CTG | (Leu/L) Leucine | L452R |
| | 7 | AGA | (Arg/R) Arginine | T19R, R158G |
| | 8 | GCC | (Ala/A) Alanine | A475H |
| | 10 | AGG | (Arg/R) Arginine | R190S |
| 5 | 1 | CTG | (Leu/L) Leucine | L452R |
| | 4 | CGT | (Arg/R) Arginine | P681R |
| | 5 | CAC | (His/H) Histidine | Q1071H, D1118H, Q677H |
| | 6 | GCC | (Ala/A) Alanine | A475H |
| | 9 | AGC | (Ser/S) Serine | R190S |
| | 10 | TTC | (Phe/F) Phenylalanine | V367F, Del157 |
| 7 | 4 | GCC | (Ala/A) Alanine | A475H |
| | 6 | ACG | (Thr/T) Threonine | K417T |
| | 7 | CGT | (Arg/R) Arginine | P681R |
| | 10 | CAG | (Gln/Q) Glutamine | Q677H |
| 3 | 3 | AGC | (Ser/S) Serine | R190S |
| | 5 | ACG | (Thr/T) Threonine | K417T |
| | 9 | CGG | (Arg/R) Arginine | L452R |
| | 10 | CAC | (His/H) Histidine | Q1071H, D1118H, Q677H |
| 15 | 1 | ACG | (Thr/T) Threonine | K417T |
| | 2 | CGG | (Arg/R) Arginine | L452R |
| | 5 | CGT | (Arg/R) Arginine | P681R |
| | 6 | AGG | (Arg/R) Arginine | R190S |
| 30 | 3 | CAT | (His/H) Histidine | H655Y, P681H, Q677H, Del 69, Q1071H |
| | 5 | CTG | (Leu/L) Leucine | L452R |

| | | | | |
|---|---|---|---|---|
| | 7 | CAC | (His/H) Histidine | Q1071H, D1118H, Q677H |
| | 8 | ACC | (Thr/T) Threonine | T20N |
| | 9 | ACG | (Thr/T) Threonine | K417T |

**Table S3.** ***Dimensions of Concern (DOCs), associated codons, and variants.***

| **Variants** | **Genomic Substitution** | **Reference codon** | **Replaced codon** |
|---|---|---|---|
| L452R | T22917G | CTG | CGG |
| K417T | A22812C | AAG | ACG |
| R190S | G22132T | AGG | AGT |
| Q1071H | A24775T | CAA | CAT |
| D1118H | G24914C | GAC | CAC |
| Q677H | G23593T | CAG | CAT |
| W152C | G22018T | TGG | TGT |
| T716I | C23709T | ACA | ATA |
| D614G | A23403G | GAT | GGT |
| H655Y | C23525T | CAT | TAT |
| P681H | C23604A | CCT | CAT |
| A570D | C23271A | GCT | GAT |
| G142D | G21987A | GGT | GAT |
| D138Y | G21974T | GAT | TAT |
| S982A | T24506G | TCA | GCA |
| V1176F | G25088T | GTT | TTT |
| T478K | C22995A | ACA | AAA |
| K417N | G22813T | AAG | AAT |
| T20N | C21621A | ACC | AAC |
| T19R | C21618G | ACA | AGA |
| E484Q | G23012C | GAA | CAA |
| E484K | G23012A | GAA | AAA |

**Table S4.** ***List of variants, corresponding genomic substitution, and associated codon change.***

| RESOURCE | SOURCE | IDENTIFIER |
|---|---|---|
| ***Data*** | | |
| SARS-CoV-2 genome sequeces | GISAID | https://www.gisaid.org/ |
| COVID-19 cases | GitHub | https://github.com/CSSEGISandData/COVID-19 |
| ***Software and algorithms*** | | |
| CoV-Seq | GitHub | https://github.com/boxiangliu/covseq |
| Flt-SNE (version 1.2.1) | GitHub | https://github.com/KlugerLab/FIt-SNE |
| Anaconda (version 4.10.1) | anaconda.com | https://www.anaconda.com/ |
| Python (version 3.8.5) | python.org | https://www.python.org/downloads/release/python-385/ |
| biopython (version 1.78) | PyPI | https://pypi.org/project/biopython/1.78/ |
| gensim (version 4.0.1) | PyPI | https://pypi.org/project/gensim/4.0.1/ |
| numpy (version 1.19.2) | PyPI | https://pypi.org/project/numpy/1.19.2/ |
| pandas (version 1.1.3) | PyPI | https://pypi.org/project/pandas/1.1.3/ |
| matplotlib (version 3.3.2) | PyPI | https://pypi.org/project/matplotlib/3.3.2/ |
| seaborn (version 0.11.0) | PyPI | https://pypi.org/project/seaborn/0.11.0/ |
| plotly (version 4.14.3) | PyPI | https://pypi.org/project/plotly/4.14.3/ |
| R version (4.1.0) | CRAN | https://cran.r-project.org/ |
| lubridate (version 1.7.0) | CRAN | https://cran.r-project.org/web/packages/lubridate/ |
| dplyr (version 1.0.7) | CRAN | https://cran.r-project.org/web/packages/dplyr/ |
| zoo (version 1.8.9) | CRAN | https://cran.r-project.org/web/packages/zoo/ |
| tseries (version 0.10.48) | CRAN | https://cran.r-project.org/web/packages/tseries/ |
| plyr (version 1.8.6) | CRAN | https://cran.r-project.org/web/packages/plyr/ |
| reshape2 (version 1.4.4) | CRAN | https://cran.r-project.org/web/packages/reshape2/ |
| ggplot2 (version 3.3.5) | CRAN | https://cran.r-project.org/web/packages/ggplot2/ |
| ggpubr ( version 0.4.0) | CRAN | https://cran.r-project.org/web/packages/ggpubr/ |
| TSEntropies (version 0.9) | CRAN | https://cran.r-project.org/web/packages/TSEntropies/ |
| DCCA (version 0.1.1) | CRAN | https://cran.r-project.org/web/packages/DCCA/ |
| Boruta (version 7.0.0) | CRAN | https://cran.r-project.org/web/packages/Boruta/ |
| randomForest (version 4.6.14) | CRAN | https://cran.r-project.org/web/packages/randomForest/ |
| ape (version 5.5.0) | CRAN | https://cran.r-project.org/web/packages/ape/ |
| iTOL (version 6.3) | Online | https://itol.embl.de/ |

**Table S5.** ***List of softwares and packages used for our study with their sources and identifiers for the reproducibility of this study.***
